# *sept-1/zina-1* is an Ancient Toxin-Antidote System in *Caenorhabditis elegans*

**DOI:** 10.1101/2025.11.28.691152

**Authors:** Dongyao Liu, Chaogu Zheng

## Abstract

The toxin-antidote (TA) systems, consisting of two tightly linked genes acting as a postzygotic distorter, are found in several eukaryotic organisms, including three androdioecious *Caenorhabditis* species. However, the evolutionary history of such TA systems remains poorly understood. We report an ancient *C. elegans* TA system identified in the highly divergent Hawaiian strain XZ1516. The maternal toxin SEPT-1 induces a rod-like larval arrest phenotype in progeny that do not carry the antidote ZINA-1, which has ten zinc finger motifs. Interestingly, *zina-1* is pseudogenized in most non-Hawaiian strains, while *sept-1* evolves from a toxic to a non-toxic form through sequence divergence. Phylogenetic studies found that *sept-1/zina-1* is the most ancestral TA system among the three known ones, and its loss preceded the rise of *sup-35/pha-1*. These two maternal TAs do not coexist in the same strain but have overlaps with the paternal TA *peel-1/zeel-1*, which is the latest one to propagate.

## INTRODUCTION

The toxin-antidote (TA) system is a type of two-component selfish genetic element that promotes its own propagation but does not contribute to the fitness of the host.^1,2^ The TA systems were first discovered in prokaryotes with the F-plasmid addition in *E. coli* as a well-known example.^3^ The plasmid carries a two-gene operon encoding a toxin CcdB that poisons the DNA gyrase and an antidote CcdA that disables the toxin; after cell division, only the daughter cells that inherit the plasmid can survive.^4^ So far, hundreds of TA elements with different mechanisms of the toxin-antidote interactions have been found in prokaryotes.^5^ In contrast, only a few TA elements are characterized in eukaryotes, including the spore killers in fungi, the gamete-killing meiotic drives in rice, the *Medea* factors in flour beetles, and the *peel-1/zeel-1* and *sup-35/pha-1* systems in nematodes.^2^

In eukaryotes that undergo sexual reproduction, the TA system often operates by transmitting the toxin into the gametes through the cytoplasm and allowing the toxin to sabotage the development of the gamete or the fertilized egg in the absence of the genomically encoded antidote, which can block the effects of the toxin.^6^ As a result, the toxin and antidote are usually coded by two closely linked genes in nearby loci, allowing them to be inherited together. In some surprising cases, they can also be coded by a single gene that exert both toxicity and resistance.^6^

The nematode *Caenorhabditis elegans* has served as an important model organism for the study of the TA systems thanks to the extensive sampling efforts in the past few decades that isolated thousands of wild isolates of *C. elegans*.^7–9^ By crossing the wild strains of *C. elegans* with the laboratory strain N2, previous studies discovered two pairs of TA elements that exist in N2 but not some wild strains. The *peel-1/zeel-1* system encodes a paternal toxin PEEL-1 which is deposited into the sperm and causes embryonic lethality if the offspring does not carry the antidote gene *zeel-1*.^10,11^ The *sup-35/pha-1* system encodes a maternal toxin SUP-35 which is deposited into the oocyte and causes defects in organogenesis and lethality unless the offspring carries the antidote *pha-1*.^12^ The toxicity and detoxification mechanisms of the two TA systems are not entirely clear, although a recent study found that PEEL-1 is likely a transmembrane cation channel that forms pores to damage the integrity of cell membrane.^13^ It is also unclear whether there are more TA elements in addition to the two known ones and whether any wild strain carries TA elements that are not present in the laboratory standard N2 strain.

*C. elegans* offers tremendous opportunities to discover and understand TA elements in animals because of the availability of numerous wild strains, the ease of doing crosses and genetic manipulation, and the rich information on gene functions through decades of molecular studies. Moreover, self-fertilization of *C. elegans* to some extent might have prevented the fixation of TA elements in the species, allowing them to be identified through the hybridization of different strains. This idea is supported by a recent study that revealed the potentially widespread presence of TA elements in other self-fertilizing *Caenorhabditis* species, such as *C. tropicalis* and *C. briggsae*.^14^

Several questions regarding the evolution of the TA systems can be addressed by identifying more TA systems. For example, what is the evolutionary process that inactivates the toxin gene? Is deletion or pseudogenization of the toxin the only way to get rid of the TA system? Can the toxin gene evolve from a toxic to a non-toxic form? For multiple TA systems, are there any patterns of ecological distribution? What is their evolutionary history? Can multiple TA systems co-exist in the same individuals or do the population switch from one to another during evolution?

In this study, we address the above questions by identifying a novel TA system (named *sept-1/zina-1*) in a highly diverged *C. elegans* wild isolate XZ1516. SEPT-1 is a maternally deposited toxin that disrupts the function of the intestine, whereas ZINA-1 is the antidote that blocks the effects of SEPT-1. Interestingly, *zina-1* is pseudogenized in N2, while *sept-1* was kept and evolved into a largely non-toxic form we named *sept-2*, which only shares 47% identity with *sept-1*. Phylogenetic analysis showed that *sept-1/zina-1* mostly exists in the Hawaiian strains that carry the most ancestral variants. This ancestral TA system disappeared entirely in the non-Hawaiian strains, and its loss coincides with the rise of another maternal TA system *sup-35/pha-1*. Such evolutionary dynamics hints at the potential incompatibility of multiple TA systems that transmit through the same route.

## RESULTS

### Genetic mapping of a new toxin-antidote element in a divergent Hawaiian strain

When crossing the laboratory standard Bristol strain N2 with a highly divergent Hawaiian strain XZ1516, we noticed that although the N2/XZ1516 hybrids (F1) developed normally, about ∼40% of their progeny (F2) from self-fertilization either died at late embryonic stages or arrested at early larval stages. Two types of larval arrest phenotypes were observed: one showed a rod-like shape, while the other showed a more curled shape with a malformed pharynx (Figure 1A). We then crossed N2 that carried different chromosomal markers with XZ1516 and found that the surviving offspring of the N2/XZ1516 hybrids must carry the chromosome III (chrIII) of N2 and chrV of XZ1516 (Figure 1B-E), suggesting the presence of two independently segregating TA systems in the hybrid. Between the two identified TA systems in *C. elegans*,^10,12^ *zeel-1/peel-1* system on chrI is conserved between N2 and XZ1516, but *sup-35/pha-1* locus on chrIII shows divergence (Figure S1A-B). In the XZ1516 genome, we found inversion and duplication events that led to sequence divergence of the toxin *sup-35* and a deletion event that likely inactivated the antidote *pha-1*. To test whether the preferred inheritance of N2 chrIII was caused by the *sup-35/pha-1* locus that was potentially nonfunctional in XZ1516, we deleted *sup-35* in N2 carrying a chrIII marker and found that the bias towards N2 chrIII disappeared among the offspring of the N2/XZ1516 heterozygotes (Figure 1F and S1C). The lethality rate dropped to ∼25% (Figure 1G) and the arrested larvae all appeared to be rod-like. These results indicated that there remained one other TA element, which was present on the chrV of XZ1516 and absent in N2. This TA system is likely new, since no chrV-associated TA was identified before.

**Figure 1.**
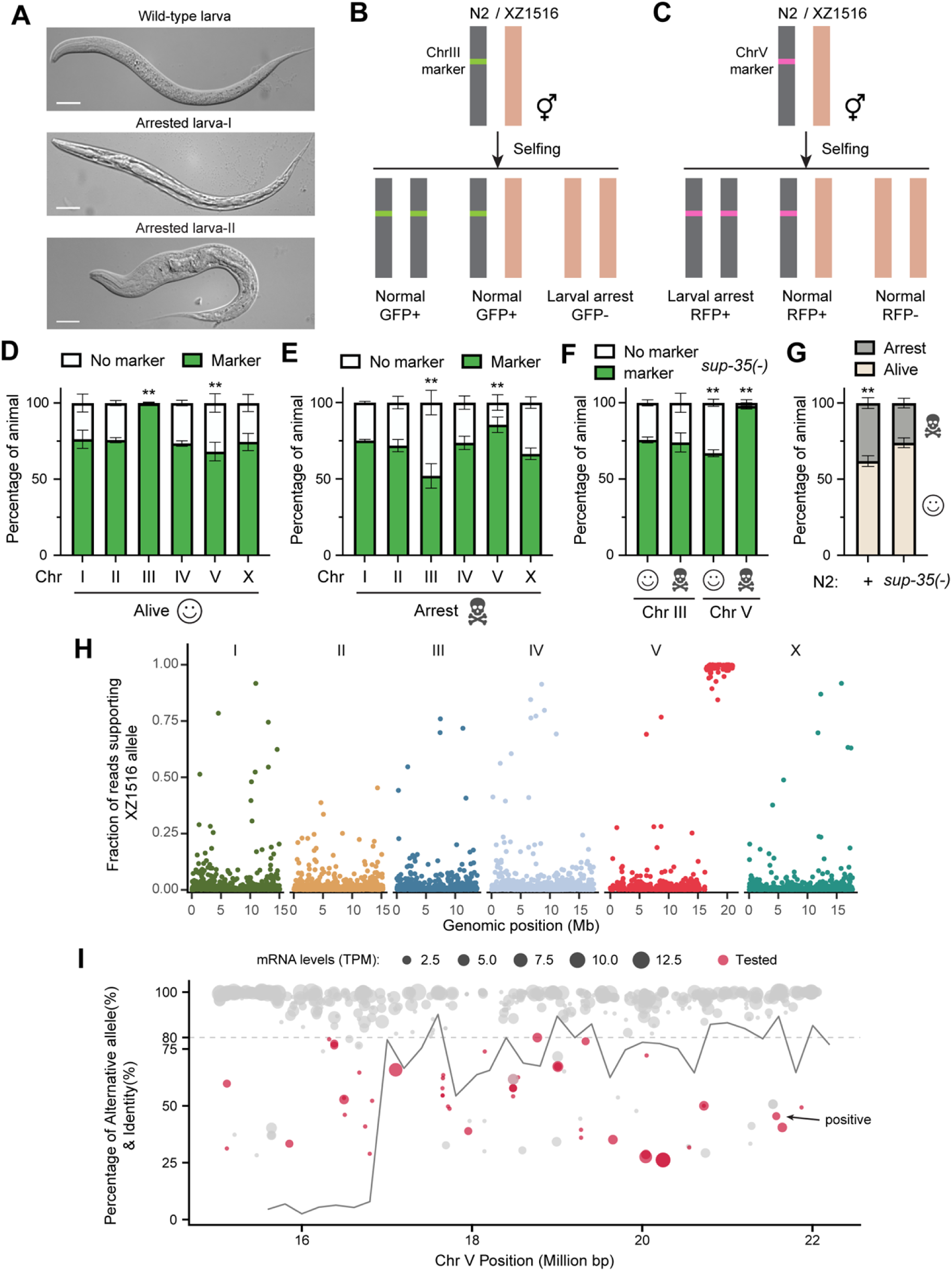
Mapping of a novel TA system in XZ1516. (A) Arrested larvae among the offspring of N2/XZ1516 hybrids. Type I represents the rod-like arrested larva, while type II represents the curled-up larva with deformed pharynx. Scale bars, 20 μm. (B-C) Schematic cartoon of the cross and phenotypes of the progeny of the N2/XZ1516 hybrids. Grey bars indicate the N2 chromosomes carrying fluorescent reporters as transgenes, and brown bars indicate the homologous XZ1516 chromosomes. (D-E) Percentages of live or arrested animals that carry the N2-derived fluorescent marker for each chromosome among the progeny of the N2/XZ1516 hybrids. (F) Percentages of live or arrested animals that carry the N2 chrIII or chrV marker among the progeny of N2[*sup-35(unk210)*]/XZ1516 hybrids. (G) Percentages of animals that are alive or arrested among the progeny of N2/XZ1516 or N2[*sup-35(unk210)*]/XZ1516 hybrids. In (D-G), double asterisks indicate statistically significant (*p* < 0.01) deviation from the expected 75% for animals carrying the marker or being alive in a Chi-square test. (H) Whole-genome sequencing results of the near isogenic lines (NILs) obtained by introgressing the XZ1516-derived TA locus into N2. Reads were mapped to a N2 genome scaffold installed with XZ1516 SNPs and small indels, and the proportion of the reads supporting XZ1516 alleles were plotted. (I) A zoom-in view of the right arm of chrV of a fully assembled XZ1516 genome. Genes in this region that code for proteins with > 100 amino acids, have detectable expression in the XZ1516 transcriptome, and are diverged (< 80% identity) from the N2 genes were marked as red dots and subjected to RNAi screen. The positive hit is indicated by the arrow. The size of the dots represents mRNA expression levels.

To understand the inheritance pattern of the new toxin, we crossed hybrid hermaphrodites with N2 males and found arrested larvae in the cross progeny, but the reciprocal cross using hybrid males and N2 hermaphrodites did not produce any arrested animals, suggesting that the toxin is maternally transmitted through the oocyte and its toxicity can be suppressed by a zygotic copy of XZ1516 chrV-linked antidote (Figure S1D-E).

To map the newly found TA system, we introgressed the TA locus into the N2 strain and generated near-isogenic lines (NILs) that carry the XZ1516-derived TA element in the N2 genomic background. Through whole-genome sequencing and SNP mapping (see the Materials and Methods for details), we mapped the TA element to a 5-Mb region on the right arm of chrV (Figure 1H). Using a chromosome-level XZ1516 genome, we predicted genes in this region and a nearby 2-Mb region (15-22 Mb in total) and found 458 genes that code for proteins longer than 100 amino acids and had detectable mRNA expression (FPKM > 0) in the transcriptome (Figure 1I). From these genes, we compiled a list of 46 candidate genes with low or no homology (< 80% identity) with N2 genes, assuming that the TA element is absent or highly diverged in the N2 strain (Table S1). We then created RNAi constructs expressing dsRNA against these candidate genes and transformed them into bacteria to allow the delivery of the dsRNA into XZ1516 strain through feeding RNAi. We expected silencing of the antidote gene to cause lethality in XZ1516 due to the unsuppressed toxicity from the toxin.

From this RNAi screen, we identified only one gene, whose knockdown led to the rod-like larval arrest phenotype similar to the arrested offspring of the N2/XZ1516 hybrid (Figure S2A). Synteny analysis showed that this gene was aligned to the N2 pseudogene *B0250.4* (Figure 2A). In XZ1516, it is a 15-exon gene encoding a 548-amino acid protein with ten C2H2-type zinc finger motifs, so we named this gene *zina-1* for “*<u>zin</u>*c finger-containing *<u>a</u>*ntidote *1*”. Sequence alignment suggests that *zina-1* is pseudogenized in N2 due to the deletion of exon 3, intron 3, and part of exon 4, leading to a frame shift and premature stops (Figure 2B). The rest exons have very high identity (>95.6%) between *zina-1* and N2 *B0250.4*, denoted as *zina-1(ψ)*, although there are still some missense variations (Figure 2B).

**Figure 2.**
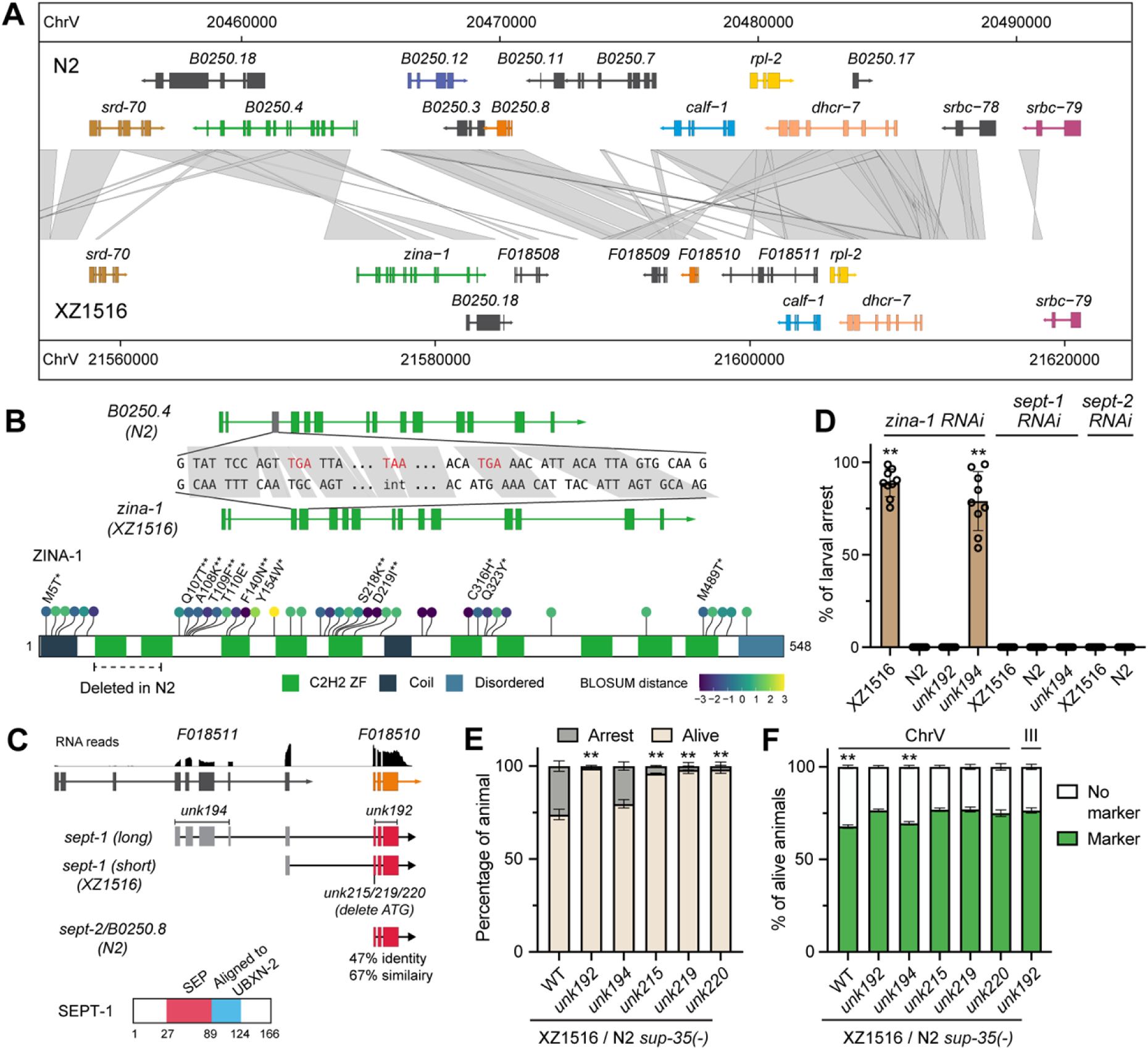
Identification of *zina-1* as the antidote gene and *sept-1* as the toxin gene. **(A)** A zoom-in view of the potential TA locus in XZ1516 and its comparison with N2 genome.*F108508*-*F108511* were *de novo* annotated by Funannotate. (B) *zina-1(XZ1516)* contains 15 exons and is pseudogenized in N2 due to the deletion of exon 3 and part of exon 4, which led to frameshift and premature stop codons (in red) from exon 3 onward. *zina-1* in XZ1516 codes for a 548 a.a. protein with ten zinc finger motifs. In addition to the deletion in N2 (dashed line), the ZINA-1 coding sequence carries many missense variants (dots with lines) in N2 compared to XZ1516. The sites that were predicted to be significantly positively selected by Bayes Empirical Bayes (BEB) analysis were labeled with asterisks. (C) RNA reads supporting the exons of *F108510* and *F108511* were shown. *sept-1* long isoform contains a 5’UTR formed by five noncoding exons and its short isoform has only one noncoding exon. It shares 47% identity and 67% similarity with B0250.8 in N2. SEPT-1 is a 166 a.a. protein with a putative SEP domain (based on similarity with *C. elegans* UBXN-2). Molecular lesions of various *unk* alleles were shown. (D) Rates of lethality caused by RNAi against *zina-1*, *sept-1*, and *sept-2* in various strains. Double asterisks indicate statistically significant (*p* < 0.01) deviation from 0% (found in XZ1516 and N2 controls) in a Chi-square test. (E) Percentages of arrested animals among the progeny of the hybrids between XZ1516 *sept-1* mutants and N2 *sup-35(-)*. (F) Percentages of live animals carrying chrIII or chrV markers among the progeny of the hybrids between XZ1516 *sept-1* mutants and N2 *sup-35(-)*. In (E-F), double asterisks indicate statistically significant (*p* < 0.01) deviation from the expected 75% for animals being alive or carrying the marker in a Chi-square test.

### Identifying a SEP domain-containing protein SEPT-1 as the toxin in the TA element

We next searched for the toxin gene based on the assumption that the toxin is closely linked to the antidote. Homology-based annotation identified *srd-70*, *B0250.18*, *calf-1*, *rpl-2*, *dhcr-7*, and *srbc-79* in the genomic region flanking *zina-1*, but they are all highly similar between N2 and XZ1516 and not likely to be the toxin. We then used Funannotate^15^ to perform *de novo* gene prediction and identified four additional protein-coding genes (*F018508 to F018511*) in this region. *F018508* had no supporting RNA-seq reads, and *F018508* showed homology with N2 *rpl-2*. *F018510* showed homology with N2 *B0250.8*, while *F018511* overlapped with *calf-1*. Since there are RNA-seq reads connecting *F018510* and *F018511*, we used forward primers that bind to spliced leaders (SL1 and SL2) and reverse primers that bind to *F018510* exon 3 to obtain full-length transcripts of *F018510* and indeed found two isoforms containing exons from *F018511* (Figure 2C). Nevertheless, these exons appeared to be noncoding and likely serve as 5’UTR. Based on RNA-seq reads, the long isoform is the minor isoform (Figure S2C).

We then deleted *F018510* coding sequence in XZ1516 and conducted RNAi against the antidote *zina-1* in the knockout mutants (*unk192*) and did not observe any arrested larvae. In contrast, deleting four exons encoding the 5’UTR of the long isoform (*unk194* allele) did not affect the lethality in *zina-1* RNAi animals (Figure 2D). These results suggest that *F018510* codes for the toxin. Since both XZ1516 *F018510* and N2 *B0250.8* code for proteins with a putative SEP domain, we named the *F018510* gene *sept-1* for “*<u>SEP</u>* domain-containing *t*oxin *1*” and the *B0250.8* gene in N2 *sept-2*, since their sequences are highly diverged and shared only 47% identity. Since the antidote is pseudogenized in N2, we hypothesize that the presumptive toxin *sept-2* may no longer be toxic in N2 due to either functional divergence or loss of expression.

To further confirm that *sept-1* is the toxin, we crossed XZ1516 *sept-1(unk192)* with N2 *sup-35(-)* and found that since both toxins are removed, the preference for N2 chrIII and XZ1516 chrV was eliminated among the offspring of the hybrids, and the fluorescent markers on both chromosomes were normally inherited by 75% of the progeny (Figure 2E-F). To further confirm that the SEPT-1 protein and not its mRNA is toxic, we created three alleles (*unk215*, *unk219*, and *unk220*) that only deleted the start codon of *sept-1* and a few flanking nucleotides (Figure S2B) and found that they indeed removed the maternal toxicity of the N2/XZ1516 hybrids (Figure 2E-F).

After mapping the TA system, we used genotyping as a more accurate method to confirm its inheritance pattern. First, in the XZ1516 background, deleting the TA on the chrV led to the loss of that chromosome in surviving offspring of the heterozygotes; the same is true for the N2/XZ1516 hybrid and the N2 backgrounds (Figure S2C and S3A). Thus, the TA element mandates its own propagation regardless of the genetic environment. Second, we confirmed that the toxin was maternally deposited into the oocytes and paternal inheritance did not cause any lethality (Figure S3B).

While we were conducting this work, the Kruglyak lab reported a TA system in XZ1516, which was discovered by crossing it with the QX1211 strain.^16^ They named the system *mll-1/smll-1*, which is likely the same as the *sept-1/zina-1* locus. However, our characterization of this TA system differs significantly (see below).

### ZINA-1 activity is lost and SEPT-2 toxicity strongly attenuated in N2

Since *zina-1(ψ)* in N2 appeared to be pseudogenized by a deletion of exon 3 and 4, we repaired the locus by inserting a 190-bp fragment of *zina-1* coding sequence to restore the open reading frame (Figure S4A). Surprisingly, this repaired allele, denoted as *zina-1(N2*)*, did not provide any protection against *sept-1* toxicity, as none of the surviving offspring of N2/XZ1516 hybrid carried the allele (Figure 3A). We also used extrachromosomal array to overexpress *zina-1(N2*)* from either the N2 *zina-1(ψ)* promoter or the XZ1516 *zina-1* promoter; neither array was able to inhibit the maternal SEPT-1 (Figure 3A-B). As controls, we expressed *zina-1* from the two promoters and both were able to protect the animals that do not carry the antidote gene in the genome (Figure 3C). This result suggests that the missense mutations in *zina-1(N2*)* may have rendered the protein inactive even if the deletion did not occur. In fact, many mutations affected residues in the zinc finger domains, potentially disrupting functions; intriguingly, several of them were positively selected (Figure 2B).

**Figure 3.**
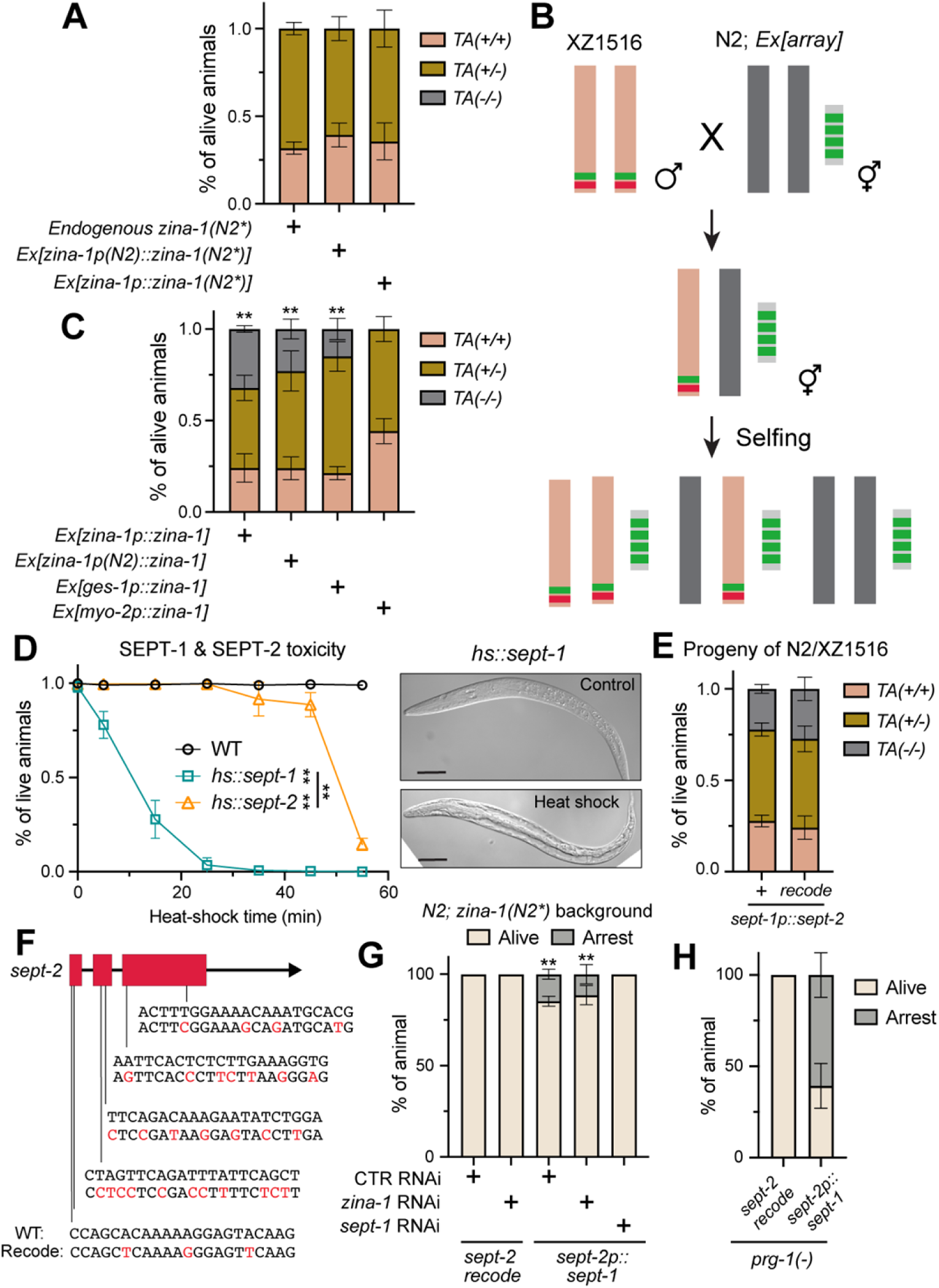
SEPT-1 proteins are more toxic than SEPT-2. (A) Percentages of live animals with specific genotypes (determined by single-worm PCR) among the progeny of N2/XZ1516 hybrid that carry the repaired *zina-1(unk232)* allele from N2, denotated as *zina-1(N2*)*, or extrachromosomal arrays expressing *zina-1(N2*)*. (B) Schematic cartoon of the experiments. Only live progeny that carried the extrachromosomal array were subjected to genotyping. (C) Percentages of live animals with specific genotypes among the progeny of N2/XZ1516 hybrid carrying the indicated extrachromosomal arrays. Double asterisks indicate statistically significant deviation (*p* < 0.01) from the expected 2:1 ratio for *TA(+/-)*:*TA(+/+)* in a Chi-square test. (D) Percentages of L1 animals (in N2 background) carrying the *unkSi32[hs::sept-1]* and *unkSi18[hs::sept-2]* that were still alive at 24 hours after a 5, 15, 25, 35, 45, and 55-min heat-shock. Representative images of *unkSi32* animals that were arrested after a 45-min heat-shock. Double asterisks indicate significant difference (*p* < 0.01) between two curves in a Kolmogorov-Smirnov test. (E) Percentages of live animals with the indicated genotypes among the progeny of N2/XZ1516 hybrid that carry the *sept-1(unk223)*, which is *sept-1p::sept-2*, or *sept-1(unk243)*, which is *sept-1p::sept-2_recode*, from XZ1516. (F) The predicted piRNA binding sites in the coding region of *sept-2*, which were mutated in the *sept-2_recode* version. Red letters indicate the synonymous mutations. (G) Percentages of animals that were arrested in N2 carrying the *sept-2(unk277, recode)* or *sept-2(unk257)* allele, which is *sept-2p::sept-1*, and subjected to various RNAi treatment. All animals carry the *zina-1(unk232*, *N2*)* background. Double asterisks indicate statistically significant (*p* < 0.01) deviation from 0% for larval arrest in a Chi-square test. (H) The same as (G) except the animals also carry *prg-1(n4503)* alleles. Details of the alleles used in this Figure can be found in Figure S4.

A puzzling issue of the *sept-1/zina-1* system is that the toxin is kept in N2 where the antidote is lost. To assess and compare the toxicity of SEPT-1 and SEPT-2 proteins, we used the MosTI approach^17^ to construct animals carrying a single-copy transgene that expressed *sept-1* or *sept-2* from a heat-shock promoter in N2 at exactly the same genomic locus (Figure S4B). Heat induction of 30 minutes produced little lethality with SEPT-2, whereas the same condition caused almost 100% lethality by SEPT-1 (Figure 3D). Although prolonged induction of SEPT-2 expression could still cause larval arrest, its toxicity was much weaker than SEPT-1 in the N2 background. To test the potential toxicity of SEPT-2 in the XZ1516 background, we swapped out *sept-1* coding region with *sept-2* sequences in the XZ1516 genome and crossed the edited XZ1516 with N2 (Figure S4C). We did not observe any lethality among the offspring of the hybrid and the edited TA elements followed the Mendelian segregation pattern (Figure 3E), suggesting that *sept-2* has no toxicity if expressed from the endogenous *sept-1* promoter. We reason that SEPT-2 has lost most of its toxicity due to protein sequence divergence, which allows *zina-1* to be pseudogenized by genetic drift.

Our findings contradict the report by Zdraljevic *et al.* from the Kruglyak lab, which suggests that *sept-2* (referred to as *B0250.8* in their paper) is not toxic because its expression in N2 is suppressed by small RNAs.^16^ To formally test this idea, we replaced the *sept-2* coding region with recoded sequences that altered most codons through synonymous mutations. This recoding likely removed the potential piRNA binding sites in the coding sequence and allowed *sept-2* to be expressed, and its toxicity be revealed (Figure 3F). However, we did not observe any larval arrest phenotype in the recoded animals (Figure 3G). Since we did the gene editing in the *zina-1(N2*)* background, we also knocked down *zina-1* and did not observe any lethality either. To further rule out the effects of piRNAs, we crossed the above animals with *prg-1(-)* mutants (*prg-1* codes for the only Piwi-family Argonaute^18^ in *C. elegans*) but still did not find any lethality caused by *sept-2* (Figure 3H). Moreover, we created animals that expressed the recoded *sept-2* from the endogenous *sept-1* promoter in XZ1516 and crossed the edited XZ1516 with N2 and did not find arrested larvae among the progeny of the N2/XZ1516 hybrids (Figure 3E and S4C). Thus, we concluded that the lack of toxicity from the *sept-2* locus is not likely due to small RNA-mediated suppression of expression but rather the non-toxic nature of the SEPT-2 protein.

As a control, we replaced the *sept-2* coding region with *sept-1* sequences and indeed observed rod-like arrest phenotype in ∼15% of the animals, supporting the idea that *sept-1* is more toxic than *sept-2* (Figure 3G). Interestingly, the lethality caused by *sept-1* expressed from the *sept-2* locus was enhanced in the *prg-1(-)* background (Figure 3H). We reason that there may be some small RNAs targeting the *sept-1* coding region or the unaltered *sept-2* UTR region in N2.

### 3’UTR of the *sept-1* mRNA has a stem-loop structure and is A-to-I edited

We next studied the expression pattern of the toxin and antidote. First, we designed single-molecule fluorescent in situ hybridization (smFISH) probes against *sept-1* mRNA but could not detect any signal in the gonad or in early embryos of XZ1516 animals. We then created an endogenous TagRFP knock-in at the *sept-1* locus in XZ1516 but failed to observe any TagRFP signals in early embryos. Antibodies staining against the HA tag fused to the C-terminus of the SEPT-1::TagRFP did not generate any signal either. Surprisingly, the smFISH against *sept-1* coding sequence gave some signals in the gonad of *sept-1::TagRFP* animals (Figure 4A). These results might suggest that the *sept-1* mRNA but not proteins are maternally deposited into the oocyte. We do not know why the same set of *sept-1* probes did not produce any signal in wild-type XZ1516 animals, but upon close inspection of the *sept-1* mRNA sequence, we discovered a very long stem-loop structure in the 795-nt 3’UTR region (Figure 4B). The entire 3’UTR of *sept-1* is mostly a palindromic sequence that folds into a hairpin structure with a 315-nt-long stem and a 7-nt unpaired loop according to a structure predicted by “RNA fold” (http://rna.tbi.univie.ac.at/cgi-bin/RNAWebSuite/RNAfold.cgi). In comparison, the coding region of *sept-1* is merely 501-nt-long. Thus, we speculate that the proximity to the hairpin-structured 3’UTR may somehow render the *sept-1* coding sequence inaccessible by the smFISH probes but inserting a 711-nt-long TagRFP sequence between the *sept-1* coding and 3’UTR regions might free up the coding region for the binding by the smFISH probes (Figure 4B).

**Figure 4.**
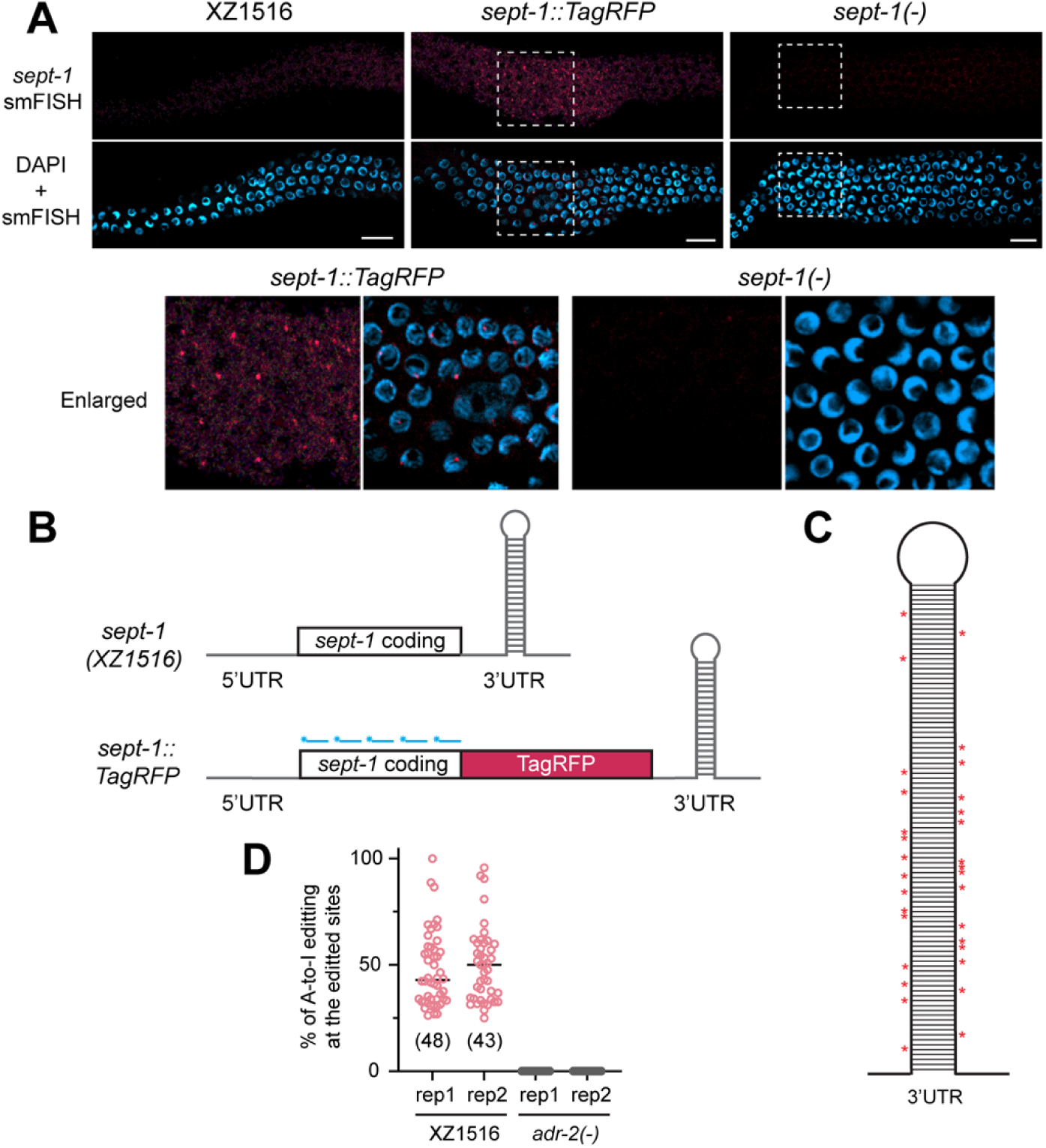
*sept-1* 3’UTR forms a hairpin structure and undergoes A-to-I editing. (A) smFISH signal against the *sept-1* coding region in dissected gonads of the wild-type, *sept-1(unk247; sept-1::TagRFP)*, and *sept-1(unk192)* XZ1516 adult animals. *sept-1(unk192)* is the *sept-1(-)* allele and serves as a negative control. The dashed boxes are enlarged in the bottom images. (B) A schematic cartoon showing the difference between the mRNAs produced by wild-type *sept-1* and *sept-1(unk247; sept-1::TagRFP)* loci. The short blue lines indicate the smFISH probes. (C) A schematic presentation of the positions of A-to-I editing in the stem structure of the *sept-1* 3’UTR based on published RNA-seq results (NCBI SRA: SRR12868314). Each asterisk indicates one edited nucleotide. The number of steps on the ladders does not accurately reflect the number of base pairings in the hairpin; the scheme is intended to show the rough positions of the edits. (D) Percentages of RNA reads that show the A-to-I editing (read as a G) for each edited site for XZ1516 wild-type and *adr-2(unk280)* animals based on the RNA-seq data generated in this study. Two biological replicates identified 48 and 43 edited sites, respectively, in XZ1516 wild-type animals; no editing was found in *adr-2* mutants.

Interestingly, when examining the published RNA-seq data (NCBI SRA: SRR12868314) of XZ1516, we found 31 “G to A” mismatches between the mRNA sequence and the genomic sequences in the *sept-1* 3’UTR (Figure 4C). Even more mismatched sites were identified from the RNA-seq data generated in this study (Figure 4D). Since inosine (I) is typically read as a guanine (G) in DNA sequencing, we concluded that the *sept-1* 3’UTR underwent extensive A-to-I editing, and this editing appeared to occur as clusters in the double-stranded region of the 3’UTR on both the ascending and descending arms of the stem (Figure 4C). Previous studies found that A-to-I editing by adenosine deaminases (ADARs) enhanced the mRNA levels of germline and neuronal genes.^19,20^ We hypothesize that the same RNA editing may help stabilize *sept-1* mRNA in the germline. To test this hypothesis, we deleted *adr-2* (the only active adenosine deaminase in *C. elegans*^20^) in XZ1516 and sequenced the transcriptomes of *adr-2(-)* mutants and found that the mismatches of RNA to DNA sequence in *sept-1* 3’UTR disappeared (Figure 4D), and the *sept-1* mRNA levels decreased by 5% (not significant). However, the loss of A-to-I editing did not affect *sept-1* toxicity, since *zina-1* RNAi in the *adr-2(-)* mutants still resulted in rod-like larval arrest and the hybrids generated by crossing *adr-2(-)* XZ1516 with *adr-2(-)* N2 still produced ∼25% progeny that were arrested. Thus, we concluded that the editing of *sept-1* RNA is not essential for its expression and toxicity. This finding is consistent with previous data in N2 that the expression level of genes whose RNAs were edited at the 3′ UTR were only slightly reduced in animals lacking *adr-1* and *adr-2* compared to the wild type.^19,20^

Using transcriptional reporters, we detected somatic expression of *sept-1* and *sept-2* in the pharynx and intestine (Figure S5). The expression level of *sept-1* appeared to be much higher than *sept-2* in both XZ1516 and N2 background, suggesting that *sept-1* promoter has stronger activity compared to *sept-2* promoter likely due to considerable sequence divergence (49.3% identity for the 2-kb promoter region). The weaker promoter may also contribute to the loss of toxicity in *sept-2*.

### ZINA-1 protects the development of the intestine by suppressing the SEPT-1 toxicity

To examine *zina-1* expression, we constructed both transcriptional and translational GFP reporters of *zina-1* in XZ1516. Widespread expression was detected in the embryos as early as gastrulation and proceeded through comma stages. The GFP::ZINA-1 translational fusion showed clear enrichment in the developing E lineage which was marked by *ges-1p::TagRFP* and formed the intestine (Figure 5A). In L1 and adult animals, we observed *zina-1p::GFP* and *zina-1p::GFP::zina-1* expression mostly in the pharynx and intestine (Figure 5B). The difference in the GFP signals between the transcriptional and translational reporters may reflect the instability of the antidote protein, which was found in other TA systems as well.^21^

**Figure 5.**
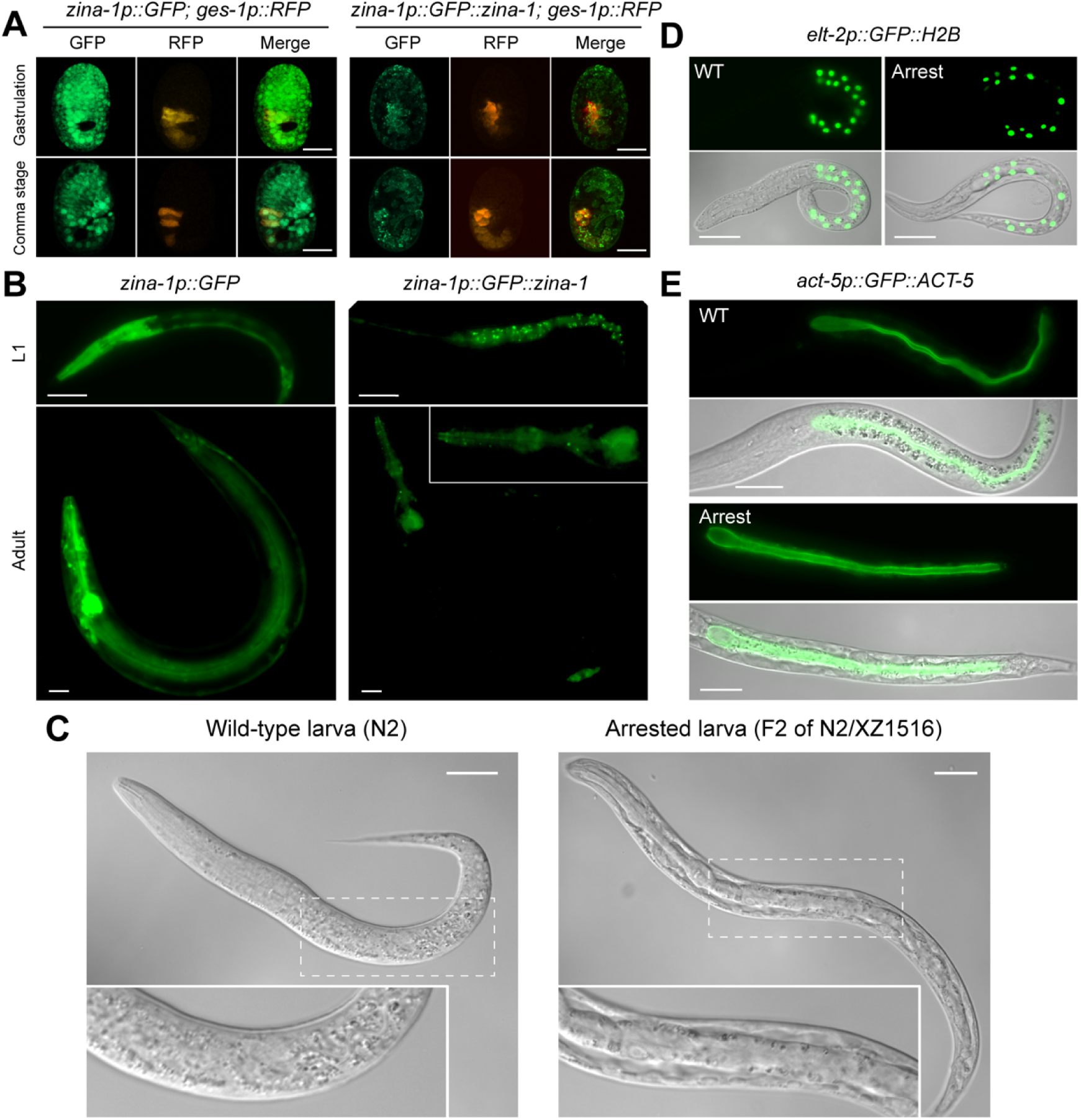
*zina-1* suppresses *sept-1* toxicity in the intestine. (A) The expression of *zina-1* transcriptional and translational reporters in the embryos at gastrulation and comma stages. The E lineage that forms the intestine is labelled by RFP expressed from *ges-1* promoter. Scale bars, 20 μm. (B) The expression of *zina-1* reporters at L1 and adult stages. The pharynx part of the adult image of *zina-1p::GFP::zina-1* is enlarged in the box. (C) Representative DIC images of a wild-type N2 larva at L1 stage and an arrested larva among the progeny of the N2/XZ1516 hybrid. Dashed boxes are enlarged in the inserts showing the dried-up body of the arrested larva. (D-E) Wild-type N2 larva and arrested larva among the F2 progeny of the N2/XZ1516 hybrid carrying the *caIs71[elt-2p::GFP::HIS-2B]* and the *jyIs13 [act-5p::GFP::act-5]* transgenes, respectively.

The enrichment of ZINA-1 in the digestive system in XZ1516 led us to hypothesize that ZINA-1 may mostly function in the intestine to detoxify SEPT-1. In fact, when we examined the arrested larva caused by the SEPT-1 toxicity, we observed abnormal intestinal morphology under the DIC; the arrested larva among the progeny of N2/XZ1516 hybrids appeared to have bloated intestine and dried up body, indicating digestion failure. Although the number of intestinal cells (labelled by *elt-2p::GFP::H2B*) were largely normal in the arrested animals, we confirmed the appearance of bloated intestinal lumen using GFP::ACT-5 that labels the apical domain of the intestinal cells (Figure 5C and 5D). These results indicated that intestinal development and/or function is disrupted by SEPT-1 in the absence of ZINA-1.

To confirm ZINA-1 functions in the intestine, we constructed extrachromosomal array that expressed *zina-1* from the intestine-specific *ges-1* promoter in N2 and crossed the animals with XZ1516. We found that if the offspring of the N2/XZ1516 hybrids carried the extrachromosomal array, they can survive even if they did not have XZ1516 chrV (Figure 3C). In contrast, pharyngeal expression of *zina-1* from the *myo-2* promoter in the extrachromosomal array could not protect the animals that did not carry the genomic *zina-1* (Figure 3C), suggesting that ZINA-1 must act in the intestine to detoxify SEPT-1.

### Ecological distribution of the three TA systems among the wild populations of *C. elegans*

The fact that N2 and XZ1516 can poison each other using two different maternal TA systems prompted us to examine the ecological distribution of the three known TA systems. We created a computational pipeline to map the presence and absence (or divergence) of the *peel-1/zeel-1*, *sup-35/pha-1*, and *sept-1/zina-1* systems among 611 wild isotypes, which were divided into three Hawaiian groups and one non-Hawaiian group based on their positions in the phylogenetic tree (Figure 6A). This strain classification is consistent with previous findings by us and others.^8,22^ Interestingly, we found that the *sept-1/zina-1* TA system was present in all strains of the “Hawaiian_1” group which carried the most ancestral variants of the *C. elegans* genome, whereas the other two TA systems were rarely found in the “Hawaiian_1” group. The frequency of *sup-35/pha-1* increased gradually from “Hawaiian_2” to “Hawaiian_3” and “non-Hawaiian” groups, while the *peel-1/zeel-1* TA system was only prevalent in the “non-Hawaiian” group. The two maternal TA systems *sept-1/zina-1* and *sup-35/pha-1* were mutually exclusive except for in one Hawaiian_1 strain; on the other hand, the paternal *peel-1/zeel-1* system co-existed with both *sup-35/pha-1* and *sept-1/zina-1* in multiple strains (Figure 6B). As expected, in many cases, we observed the loss or divergence of the toxin while the antidote is preserved. The above results led to a model that *sept-1/zina-1* is the most ancestral TA system that was gradually lost and replaced by *sup-35/pha-1* during the evolution of *C. elegans*, and *peel-1/zeel-1* may be the most recently emerged TA system or at least only propagated in the population outside of Hawaii in recent evolution.

**Figure 6.**
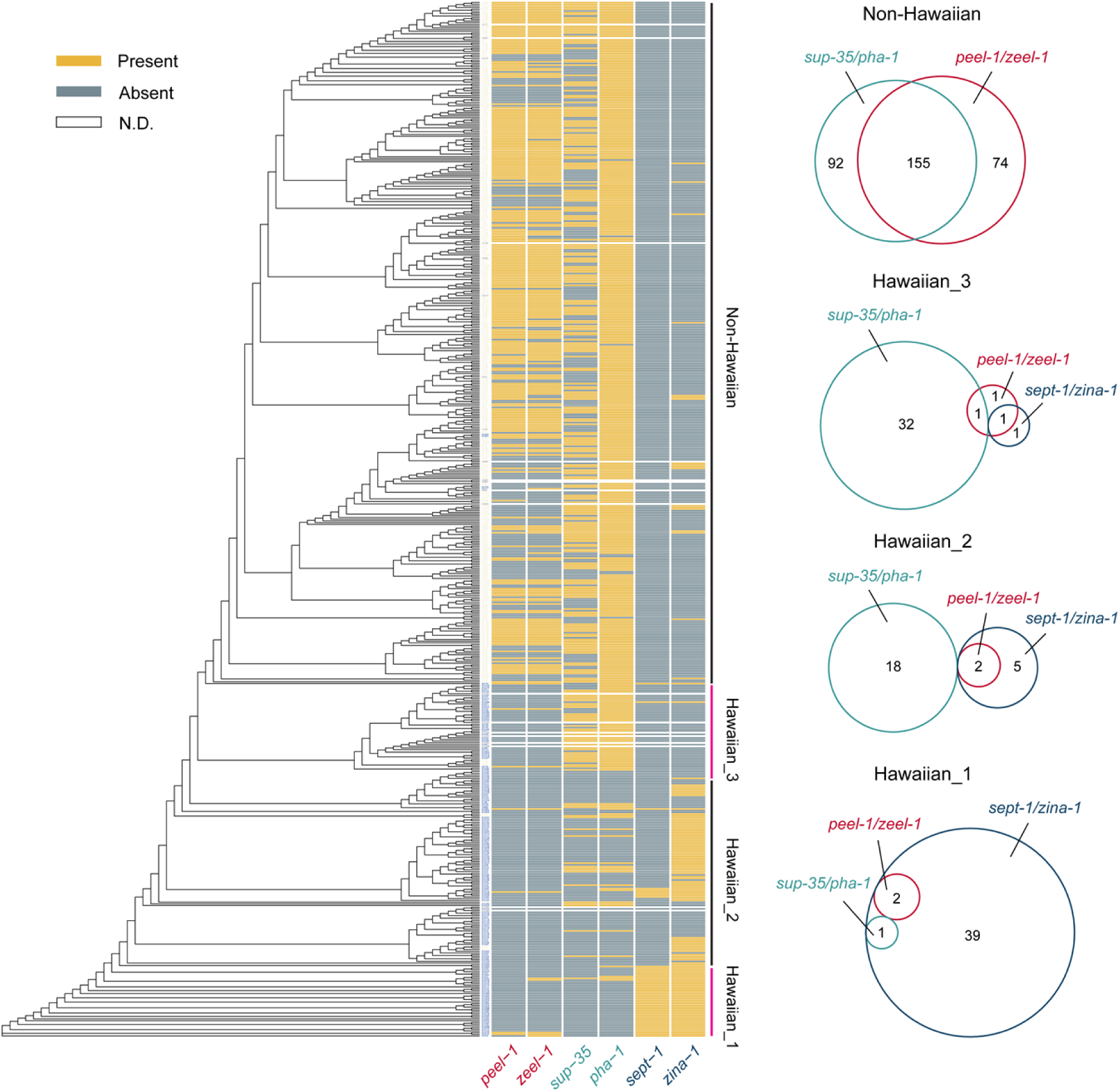
Evolutionary history of the three TA systems in *C. elegans*. A phylogenetic tree among the 611 *C. elegans* wild isotypes constructed using optimal comminated bayes models selected by ModelFinder.^32^ The strains were divided into four groups based on their clustering in the tree. Strains that were physically isolated from Hawaiian Islands are labeled in blue. The genotypes for both the toxin and antidote in the three TA systems were annotated. Sequences similar to *peel-1/zeel-1* and *sup-35/pha-1* in N2 and *sept-1/zina-1* in XZ1516 were annotated as “present”, while deletion and pseudogenization were annotated as “absent”. Sequence divergence of the toxin was also annotated as “absent”. For a few strains, we could not reliably determine the genotype of the TA systems from the whole-genome sequencing data and were labeled as “N.D.”. The overlapping among the three functional TA systems in each group of strains were shown in the Venn diagrams on the right. The number of strains that carry the corresponding TA system is shown.

It is worth noting that previous work suggested that *pha-1* was deleted and *sup-35* was pseudogenized in the DL238 strain leading to the maternal-effect incompatibility with the N2 strain. Upon close inspection of the assembled genome of DL238, we found an open reading frame for *sup-35* and the predicted SUP-35^DL238^ proteins showed significant sequence divergence from the SUP-35^N2^ protein (Figure S6). Similar divergence in SUP-35 was found in other strains including XZ1516 and likely led to the loss of SUP-35 toxicity. This result mirrors the effect of sequence divergence between SEPT-1^XZ1516^ and SEPT-2^N2^ and led us to conclude that the inactivation of a TA system does not require the deletion or pseudogenization of the toxin. Accumulation of missense mutations in the toxin gene may be a common way to disable its toxicity while potentially maintaining its other functions.

### The loss of a TA system correlates with large genetic diversity in the linked region

Given the genetic variation and evolutionary dynamics of the *sept-1/zina-1* and *sup-35/pha-1* loci in the Hawaiian groups, we divided the Hawaiian strains into two genotypes including *sept-1(+)*, representing the *sept-1(XZ1516)* allele, and *sept-1(-)*, representing the *sept-2(N2)* allele or the deletion of the *sept-1* gene. Similarly, we also divided the strains into *sup-35(+)*, representing the *sup-35(N2)* allele and *sup-35(-)*, representing the diverged or deleted *sup-35*. We found that the presence of functional *sup-35(+)* allele was correlated with lower genetic diversity on the chrIII region that harbors the gene, while the *sept-1(+)* allele was also correlated with lower polymorphisms of the nearby region on chrV (Figure 7). These finding suggests that the presence of the TA system may lead to selective sweep that reduces diversity while the inactivation of the TA system allows the accumulation of variants in the genomic region. Supporting this idea, we found more negative Tajima’s D in the region linked to *sept-1(+)* or *sup-35(+)* compared to the same region in strains without the TA system (Figure 7). Thus, the evolution of TA systems may help shape the genomic landscape of genetic variants and contribute to selective sweep.

**Figure 7.**
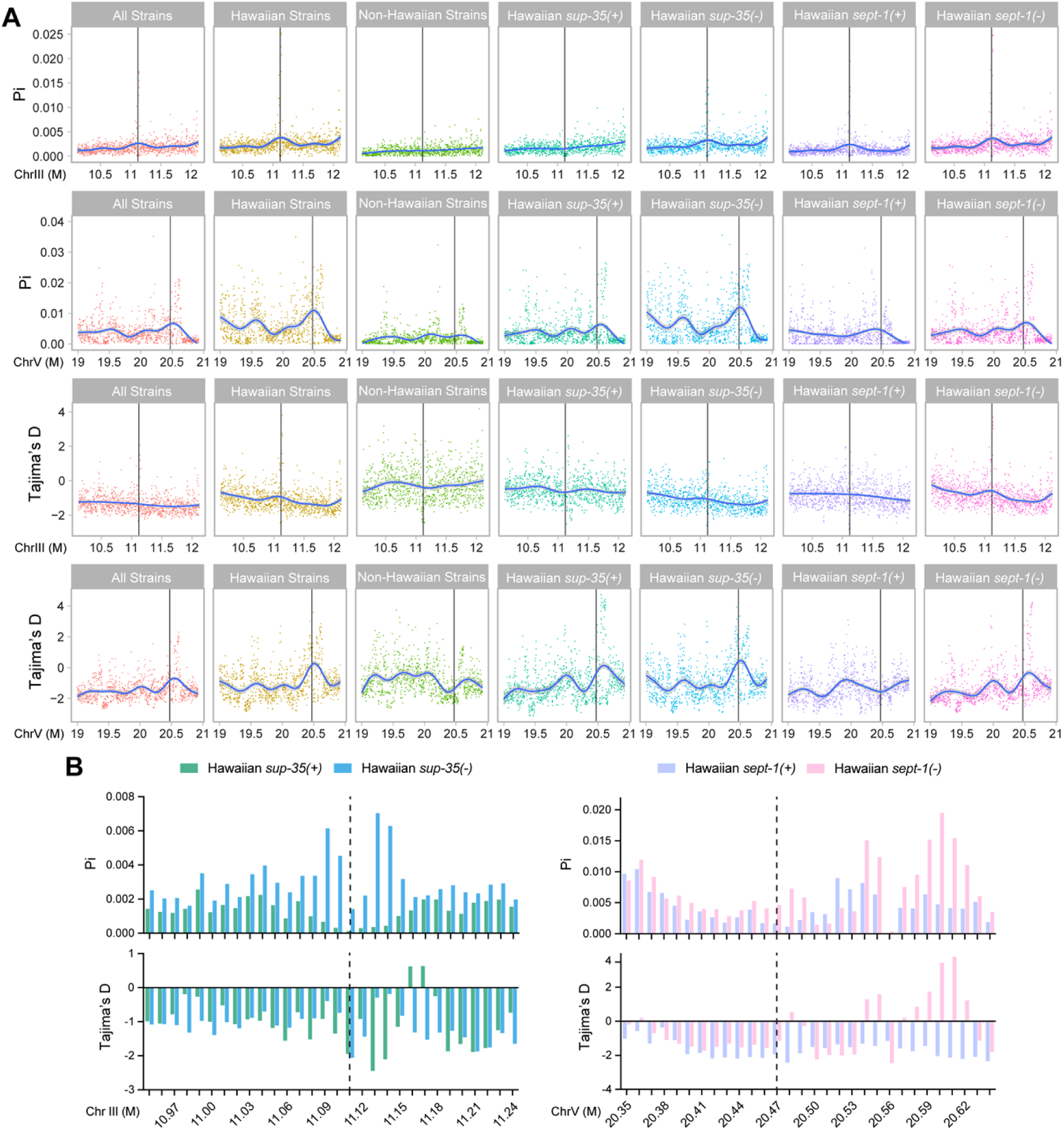
The presence of TA system may facilitate selective sweep of nearby genomic regions. (A) A 2-Mb region surrounding the *sup-35/pha-1* and *sept-1/zina-1* were analyzed for nucleotide polymorphism (Pi) and selective pressure (Tajima’s D) among all C. elegans wild strains, Hawaiian strains (including Hawaiian_1, Hawaiian_2, and Hawaiian_3 groups), and non-Hawaiian strains, as well as Hawaiians strains that carry or do carry the TA system. The black bars indicate the position of the *sup-35/pha-1* locus on chrIII and *sept-1/zina-1* locus on chrV. The nucleotides were binned into 2kb-long windows; the value for each window is shown as a dot, and the curve is calculated by generalized additive models using the R package “mgcv”. (B) The Zoom-in view of a 300-kb region centered around the *sup-35/pha-1* and *sept-1/zina-1* (positions are indicated by the dashed line). Nucleotides were binned into 10-kb windows, and the Pi and Tajima’s D values for the Hawaiian strains carry the TA or not were shown as bars.

## DISCUSSION

In this study, by crossing the laboratory N2 strain and the highly diverged Hawaiian XZ1516 strain, we identified a new TA system in the XZ1516 and named it *sept-1/zina-1*. This TA system is likely the same as the *mll-1/smll-1* system identified by Zdraljevic et al., in a recent study that crossed XZ1516 with QX1211 strain.^16^ We and their studies independently mapped the TA system to the same region on the right arm of the XZ1516 genome and showed evidence that the toxin is maternally transmitted into the oocyte likely through the deposit of toxin mRNA. We and their studies both found that the toxin has a homolog *B0250.4* (we named it *sept-2*) in the N2 strain, while the antidote is pseudogenized. However, our explanation of this unusual evolutionary pattern differs significantly.

The Zdraljevic *et al.* study suggests that *sept-2* is not toxic because its expression is suppressed by 22G siRNAs that target the piRNA binding sites in *sept-2*.^16^ In contrast, our work indicate that the toxicity of the SEPT-2 is significantly attenuated by protein sequence divergence to the point that the presence of the antidote is no longer required for the survival of the animals in the N2 strain. Even when the small RNA-mediated suppression of the *sept-2* is removed through recoding the sequence or mutating *prg-1*, we still did not observe any rod-like arrested larva. Moreover, when expressed from the *sept-1* locus in XZ1516 background, *sept-2* did not cause any lethality either in its original or recoded sequences, supporting that *sept-2* protein is nontoxic. Our findings suggest that even if *sept-2* (or *B0250.8*) is targeted by the endogenous RNAi pathway, the suppression of its mRNAs is not likely the major reason for the lack of toxicity in the absence of the antidote. Instead, the divergence of the actual protein sequences (only 47% identity) between SEPT-1 and SEPT-1 counts for most of the differences in toxicity. Interestingly, we suspect a similar coding sequence drift led to the loss of toxicity in the SUP-35 proteins in DL238, XZ1516, and other strains compared to its toxic form in N2.

In addition, the promoter of *sept-1* also appears to be stronger than *sept-2* promoter, and the stem-loop structure in *sept-1* 3’UTR and its A-to-I editing further stabilizes *sept-1* mRNA in the germline. On the contrary, *sept-2* does not have a similar 3’UTR. Thus, we concluded that multiple mechanisms enabled the evolution of the *sept-1* locus from a toxic allele in XZ1516 into a non-toxic allele in N2.

Although the function of SEPT-1 and how it poisons the intestine are unknown, the antidote ZINA-1 has ten clearly defined zinc finger motifs. Since ZINA-1 is not localized in the nucleus, we suspect that it may not be a DNA-binding protein and is instead an RNA-binding protein. One appealing hypothesis is that ZINA-1 may block the effects of SEPT-1 by binding to the *sept-1* mRNA. The long hairpin structure of the *sept-1* 3’UTR may be a potential binding site for ZINA-1. Intriguingly, the GFP::ZINA-1 appear to form aggregates both in the developing intestine and in the pharynx. This observation suggests a plausible detoxification mechanism that involves liquid-liquid phase separation, since many RNA-binding proteins were known to form condensates that help prevent the translation of the associated mRNAs.^23^

Our phylogenetic analysis uncovered an evolutionary history of the three known TA systems in *C. elegans*. We found that the *sept-1/zina-1* is the oldest TA that exists in the Hawaiian_1 strains carrying the most ancestral variants of the species. Hawaiian_2 strains represent a transition period when the *sept-1/zina-1* was gradually lost through the accumulation of missense mutations in the *sept-1* coding sequences while the antidote *zina-1* was kept and not pseudogenized yet. Coinciding with the loss of *sept-1/zina-1*, another maternal TA *sup-35/pha-1* emerged and started to propagate within the Hawaiian_2 population. Interestingly, majority (84/111; 75.7%) of the strains within this population does not carry any of the three TA system (only considering the existence of the toxin). Among the more derived Hawaiian_3 strains, the frequency of *sup-35/pha-1* increased dramatically and became the dominant TA system. The paternal TA *peel-1/zeel-1* only became prevalent in the most recently evolved non-Hawaiian population, indicating that its propagation within the species occur the latest among the three TA systems. Importantly, *peel-1/zeel-1* emerged early in the Hawaiian population but somehow did not propagate to high frequency until later in evolution.

Another interesting observation is that the two maternal TAs are almost entirely exclusive to each other, and only one Hawaiian_1 strain carried both *sept-1/zina-1* and *sup-35/pha-1*. This finding suggests a potential energy cost of maintaining two TA systems that both transmit through the oocytes. Selection may favor keeping only one. In contrast, we saw considerable overlapping of the paternal TA *peel-1/zeel-1* with either of the two maternal TA systems, indicating that TA systems may co-exist if their transmission routes are different.

## ACKNOWLEDGMENTS

We than Miss Yanjun Gao for technical assistance in genotyping some animals. We thank CaeNDR for sharing *C. elegans* wild isolate strains and their genomic data. This work is supported by the Research Grants Council of Hong Kong (GRF 17107325, GRF 17113324, and GRF 17105523). Some strains were provided by the Caenorhabditis Genetics Center, which is funded by the National Institutes of Health (NIH) Office of Research Infrastructure Programs (P40 OD010440). Computational works were performed using research computing facilities offered by Information Technology Services at the University of Hong Kong.

## AUTHOR CONTRIBUTIONS

C.Z. and D.L. conceived the study and wrote the manuscript. C.Z. initially observed the arrested phenotype among the offspring of N2/XZ1516 hybrids. D.L. carried out all the other experiments and analyzed the data. C.Z. secured funding and supervised the project.

## MATERIALS AND METHODS

C. elegans strains

*C. elega*ns strains were maintained at 20°C on nematode growth medium (NGM) plates seeded with OP50 bacteria according to previous methods.^24^ The XZ1516 strain was requested from the CaeNDR database.^9^ Transgenesis was performed by injecting the recombinant DNA into the gonad of young adults, and transgenic animals were maintained by picking the transformants with fluorescent co-injection markers. For CRISPR/Cas9-mediated gene editing in *C. elegans*, the recombinant Cas9, sgRNA, and the repair template were injected together into the worm gonads, and the progeny were screened by PCR-based genotyping. Some strains were provided by the *Caenorhabditis* Genetics Centre (CGC). All strains used in this study are listed in the Table S2.

### Alignment, variant calling, genome assembly, gene annotation, and visualization

In general, the alignment of the short-read sequencing results was done using STAR (v2.7.11b) and bwa mem (v0.7.19-r1273) with default parameter for splicing and non-splicing alignment, respectively. Deepvariant (v1.5.0) were used for variant calling. Lastz (v1.04.15) and mummer (v3.0.0) were used for large scaffolds pairwise alignment. Bedtools (v2.31.1), Conda (v24.11.3), Singularity (v3.8.0), and snakemake (v7.25.4) were used to coordinate computational tasks, environment and reproducibility on managed high-performance cluster. The scaffold-level genomes of *C. elegans* wild isolates were download from NCBI [PRJNA692613], ragtag (v2.1.0) was used to assemble the wild isolates genomes into chromosome level with the guide of N2 reference genome (WS235). Funannotate (v1.83) was used to do *de novo* annotate the chromosome-level genomes. Blastp (v2.16.0+) was used to compare the homology of proteome between wild isolates and N2. Exonerate (software, v2.80.0) was used to re-annotate the conversed or pseudogenized gene in wild isolates.

Visualization of the genomic data is performed in R using ggplot2(v3.5.2), ggtree(v3.12.0), and a self-built R package ggexon (https://github.com/DongyaoLiu/ggexon). The omega ratio of each amino acid site of ZINA-1/ZINA-1(N2*) were calculated by PAML (4.10.9) codeml model with default parameters. Software used in this study are listed in Table S3. All original code has been deposited at the github repository (https://github.com/DongyaoLiu/ToxinAntidote).

### Chromosomal mapping of the TA in XZ1516

For chromosomal mapping, we used the transgenes *zdIs5[mec-4p::GFP] I*, *muIs32[mec-7p::GFP] II*, *uIs31[mec-17p::GFP] III*, *uIs115[mec-17p::TagRFP] IV*, *uIs134[mec-17p::TagRFP] V*, and *uIs130[lad-2p::GFP] X* as the chromosomal markers for N2 and counted the percentages of live and arrested animals carrying the fluorescent marker among the offspring of the N2/XZ1516 hybrid. At least three biological replicates were carried out, and more than 500 F2 progeny were examined for each marker.

### Introgression and the mapping of the TA elements

To introgress the XZ1516-derived TA system into the N2 strain, we crossed N2 males with XZ1516 hermaphrodites and picked N2/XZ1516 hermaphrodites and cross them with N2 males again. We then repeated the cross for seven rounds and obtained the near isogenetic line named CGZ1589. To make sure CGZ1589 carries the TA system, we cross it with N2 and found rod-like arrested animals among the progeny of N2/CZ1589 hybrids. We then extracted genomic DNA from CGZ1589 using the Puregene tissue core Kit (Qiagen, cat. 1126829). 10 μg of the DNA was used for library construction and sequencing by BGI (Beijing Genome Institute, China) on the DNBSEQ PE150 platform. The sequencing depth is ∼100X. Raw sequencing file of the introgressed strain is uploaded to NCBI Sequence Read Archive (SRA) under the accession number PRJNA1359096.

To generate high-quality genomic variants between XZ1516 and N2. We use ART (v2016-06-05) to simulate 200bp pair-end reads from N2 and XZ1516 (with the 50x depth) and map the simulated reads to each other genome. We used deepvariant (v1.5.0) to identify N2 variants based on the XZ1516 genome and XZ1516 variants based on the N2 genome and then intersected the two lists of variants to identify the final high-quality variants. We then mapped the sequencing reads of the introgressed strain CGZ1589 to the N2 reference genome and counted the reads that support either the N2 variants or XZ1516 variants (at the high-quality variant sites with alternative alleles) across the genome. The read depth of different alleles of CGZ1589 were quantified using bam-readcount (v1.0.1). The region that contains most reads supporting the XZ1516 variants was identified as the TA-containing region (17-22 Mb on the right arm of chrV).

We then filtered the genes annotated in this region of the XZ1516 genome for gene size (coding proteins longer than 100 amino acids), expression levels (with supporting RNA-seq reads), and the level of homology with N2 genes (lower than 80% identity) and obtained a list of genes for RNAi screen. To include the possibility that the TA system may be located on the scaffolds that were not assembled into the chromosome-level XZ1516 genome, we also used Augustus (v3.5.0) to predict the genes located on any scaffolds aligned to N2 chrV and identified genes with the support of RNA-seq reads (PRJNA669810).

### RNAi screen for the antidote gene

Feeding RNAi was carried out using a previously described protocol.^25^ The cDNA of the 46 candidates for the antidote gene were cloned from the cDNA library of XZ1516 and inserted between the two T7 promoters in the L4440 vector using Gibson Assembly (primers used for the cloning were listed in Table S1). The resulted DNA constructs were transformed into *E. coli* HT115 strain. The bacteria containing the RNAi constructs were seeded onto NGM plates with 5 mM IPTF to induce the expression of dsRNAs. The next day, 30 bleach-synchronized embryos were placed onto the RNAi plates. The next generation were examined for the presence of rod-like arrested larvae.

### Genotyping for the inheritance pattern of the *sept-1/zina-1* locus

Live F2 progeny generated by various F1 hybrids were subjected to single-worm PCR. For the experiments that involves extrachromosomal arrays expressing the antidote, we only genotyped F2 animals that were alive and carried the array. At least three replicates were carried out, and more than 96 animals were genotyped for each cross. The genotyping primers were listed in Table S4.

### CRISPR/Cas9-mediated gene editing

Gene editing was conducted using our previously published protocol.^26^ Cas9 cut sites were predicted using CHOPCHOP (https://chopchop.cbu.uib.no). Single-guide RNAs were prepared by *in vitro* transcription using the NEB sgRNA synthesis kit (E3322S) and were incubated with recombinant SpCas9 protein (NEB # M0646T) at 37°C for 15 minutes. The RNP was then mixed with a linearized muscle-expressing co-injection marker construct (CGZ#400, *ttr-16p::GFP*), repair template (if needed), and filler DNA. For insertion or substutition longer than 150 bp, we used a previously described PCR method to generate repair template with overhangs.^27^ For edits shorter than 150 bp, single-stranded donor oligos were used as repair template. After the injection, transformants were picked to individual plates and genotyped by PCR and Sanger sequencing to find the correctly edited allele. In some cases, we also used a co-CRISPR protocol^28^ to screen for the successfully edited animals. The sequences of the CRISPR/Cas9 target and repair templates used to generate the edited alleles were listed in Table S4.

### DNA constructs and expression reporters

To generate the transcriptional reporters, a 2-kb *sept-1* promoter and a 2-kb *sept-2* promoter were cloned from the genomic DNA of XZ1516 and N2 strains, respectively, and then ligated upstream of TagRFP and a *tbb-2* 3’UTR in the pUC57 backbone. For the *zina-1* rescue experiment, *zina-1* fragment containing a 2-kb *zina-1* promoter, the entire coding region, and a 1-kb *zina-1* 3’UTR was cloned from XZ1516 genomic DNA. We then swapped out the *zina-1* promoter with a 2-kb promoter upstream of the *zina-1(ψ)*, a 2-kb *ges-1* promoter, or a 2-kb *myo-2* promoter from the N2 genomic DNA. To test the detoxification ability of *zina-1(N2*)*, which is the repaired *zina-1* allele in N2, we cloned the *zina-1* coding sequence from the cDNA library of animals carrying the *zina-1(unk232)* allele or substituted the exon 3 of the *zina-1(ψ)* cloned from N2 cDNA library with the exon3 and exon4 of *zina-1* from XZ1516 cDNA library. This *zina-1(N2*)* fragment was then placed downstream of either *zina-1* promoter or the *zina-1(ψ)* promoter. To make *zina-1p::GFP* transcriptional reporters, a 2-kb *zina-1* promoter and a 1-kb *zina-1* 3’UTR were cloned from the XZ1516 genomic DNA and were ligated to the upstream and downstream of GFP coding sequences, respectively. To make *zina-1* translational reporter, GFP coding sequence is inserted upstream of the *zina-1* coding sequence in the *zina-1* fragment containing the promoter, the coding region, and the 3’UTR cloned from the XZ1516 genomic DNA. All DNA constructs generated and used in this study are listed in Table S5.

### smFISH, antibody staining, and microscopy imaging

smFISH probes against the *sept-1* coding region were designed using the Stellaris RNA FISH Probe Designer tool from Biosearch Technologies (Petaluma, CA) and synthesized them with Quasar 670 dye. Dissected adult gonads and early embryos from various strains were fixed, stained with the FISH Probe sets in the hybridization buffer (purchased from Biosearch Technologies), and then washed using the wash buffer based on a previous protocol.^29^ For antibody staining, we used the Ruvkun protocol.^30^ Early embryos fixed in the fixation buffer and the underwent three rounds of freeze-thaw cycles. Anti-HA Mouse Monoclonal Antibody (TransGene, HT301-02) and anti-mouse secondary antibodies (Thermo-Fisher, #31430) were used. Lecia fluorescent microscopy (DMi8 with K5 monochrome camera) was used to image the larvae and adults. Embryos and dissected gonads were imaged on Confocal Laser Scanning Microscope (Carl Zeiss LSM 980).

### RNA sequencing and mapping of A-to-I editing

Mix-stage worms were collected from NGM plate. Two biological replicates were prepared for each stain.1 mg of total RNA were used for beads-based mRNA capturing and library construction and were sequenced on DNBSEQ-T7 by BGI. 6G adaptor-trimmed reads were used for downstream analysis. All raw RNA-seq data were uploaded to NCBI SRA under the accession number PRJNA1359096. Mapping followed the protocol mentioned above. The A-to-I mutation were identified by bcftools (v1.21).

### Phylogenetic tree and mapping TA systems in wild isolates

The phylogenetic tree for the wild isolates was calculated by iqtree(v1.5.5) using concatenated CDS of 4,736 single-copy gene defined by previous studies.^31^ The CDS variations were extracted from vcf file and install to the reference transcripts using a modified script (https://github.com/DongyaoLiu/vcf2fasta). Genomic data of *C. elegans* wild isotypes and the corresponding hard-filtered vcf file were downloaded from CaeNDR (v20231213).

Fastq file were extract from the bam alignment file. To map *peel-1*/*zeel-1* and *sup-35*/*pha-1* in *C. elegans* wild isolates. we combined the information from variant annotation generated by ANNOVAR (v) and reads depth at the gene location. For the new identified *sept-1*/*zina-1* and *sup-35^#^(DL238)*, we extracted the unmapped reads by samtools (v1.16.1) from the bam files and mapped those reads to XZ1516 and DL238 genomes. We called the variants within the region V: 21566743-21626152 in XZ1516, and the region III: 11247407-11301373 in DL238 and then annotated the variants. For all three TA systems, if we found nonsense mutation in the coding region or if there were no reads mapped to the region, we consider the gene to be pseudogenized. For *sept-1*, genotypes matching *sept-2(N2)* were considered as *sept-1(-)*. For *sup-35*, genotypes matching *sup-35^#^*^1^*(XZ1516)*, *sup-35^#^*^2^*(XZ1516)*, or *sup-35^#^(DL238)* were all considered as *sup-35(-)*.

### Population genetics studies

R package Genopop (v1.0.0) was used to calculate *Pi* and Tajima’s *D* from the hard-filtered vcf variant files. Two binning conditions (2 kb and 10 kb) were used for the calculation at different genomic resolution. The population groups were constructed based on the phylogenetic tree, as well as the genotype for the TA loci.

### Statistical analysis

All quantitative data were presented as mean ± SD. At least three biological replicates were performed. For categorical data (percentage of animals carrying the marker or being alive), we used Chi-square test to detect the statistically significant deviation from the expected frequencies. To compare two curves, we used the non-parametric Kolmogorov-Smirnov (KS) test. Detailed statistical information can be found in the figure legends.

